# An adverse rearing environment alters maternal responsiveness to infant ultrasonic vocalizations

**DOI:** 10.1101/2024.03.26.586776

**Authors:** Alekhya K. Rekapalli, Isabel C. Roman, Heather C. Brenhouse, Caitlyn R. Cody

## Abstract

Rodent pups use a variety of ultrasonic vocalizations (USVs) to facilitate maternal care. Importantly, infant USV repertoires are dependent on both the age and early life experiences of the pups. We have shown that an adverse rearing environment modeled with the maternal separation (MS) paradigm alters caregiving behavior, but little is known about how pup USVs differentially elicit maternal attention. In the present study, maternal approach towards a vocalizing pup over a non-vocalizing pup was tested in a Y-maze apparatus at two developmental time points over the course of MS. At the postnatal day (P) 10, dams engaged in longer interaction times with the vocalizing pup compared to the non-vocalizing pup. This effect was modulated by rearing environment and the sex of the pup, with only MS dams spending more time with vocalizing male pups. As expected at P20, dams did not show a preference for either the vocalizing or non-vocalizing pups regardless of rearing environment, however, MS dams spent a greater amount of time in the center of the apparatus as compared to control dams, which can be interpreted as a measure of uncertainty and indecision. These effects are important considering the sex specific effects of MS exposure across all developmental stages. Our novel findings support the hypothesis that sex-specific pup-dam interactions may drive later life outcomes following adversity.

## Introduction

The caregiver-offspring relationship is the first and most important relationship an altricial infant will form in the early stages of life. This early postnatal period encompasses drastic growth, neural pruning, and development, which is dependent on the bond formed between the caregiver (the mother in most species) and infant (Winston and Chicot, 2016). This relationship is defined by a series of transactions, in which one organism emits a cue, and the other responds and adapts their behavior to suit the needs of the dyad (Jenkins et al., 2016). Rat pups are born with limited mobility, and therefore rely heavily on ultrasonically vocalized (USV) communication to obtain their needs from the dam (Muller et al., 2010, Branchi et al., 2001; Ehret, 2005; Wöhr and Schwarting, 2008; Lingle et al., 2012). Rats use USVs to communicate both appetitive and aversive signals throughout their lifespan (Knutson et al. 2002; Brudzynski 2005; Burgdorf et al., 2008; Brudzynski 2009; Whör & Schwarting 2013; Brudzynski 2013), and the development of rodent USV communication patterns can provide insight into the trajectory of pup maturation.

Early life adversity (ELA) is known to cause altered maturation, which has been shown to reroute brain development (Gee et al., 2013; Honeycutt et al., 2020), disrupt performance on cognitive and learning tasks (Grassi-Oliveria et al., 2016, Goodwill et al., 2018, Bath et al., 2017), cause HPA dysregulation and immune system dysfunction (Coley et al. 2019, Bolton et al., 2018; Lehmann et al., 2002; Aisa et al., 2007; Enoch, 2011; Chocyk et al., 2013; Wieck et al., 2013; Holland et al., 2014), and alter weight gain and reflex development (Demaestri et al., 2020). Adverse rearing environments have also been shown to decrease predictability of maternal care (Granata et al., 2022a, Demaestri et al., 2022) and increase abuse-like behavior (Gallo et al., 2019). In contrast, offspring of attentive and consistent caregiving show fewer anxiety symptoms, lower levels of fearfulness, and heightened trans-generational care to their own offspring (Caldji et al.,1998; Champagne et al., 2001; Francis & Kuhar, 2008). Thus, various groups have investigated the impact of the home cage environment on the mother-infant relationship. However, a challenge remains to understand how mothers respond directly to pup communication based on rearing environment or individual pup characteristics.

In addition to the myriad cognitive and systemic effects of ELA on pup development, prior work has shown that the developmental trajectory of USV patterns is altered by ELA exposure, and that the specific USV parameters altered by ELA are sex and adversity dependent (Granata et al. 2021). Maternal Separation (MS) is a paradigm of ELA where rat pups are separated from dams for 3-4 hours per day during the pre-weaning period. MS alters the development of USV properties, but only in males; for example, the number of calls emitted in response to a brief isolation increases between postnatal day (P)5-10 for all pups, yet MS exacerbates this developmental increase, such that at P10, MS males vocalize significantly more than control-reared males. Altered developmental trajectories of USVs can be predictive of social behaviors in adolescence (Granata et al. 2022a), and therefore can be viewed as an important measure of psychological development. However, while differential USV development and maternal care individually lend insight into altered maturation, the relationship between altered USV communication and the mother-pup dyad is elusive. Therefore, in an exploratory analysis we aimed here to determine whether maternal responsiveness to these previously identified USV changes was altered during rearing adversity, and whether responsiveness differed as a function of the sex of the pup.

## Methods

### Animal husbandry procedures

All experiments were performed in accordance with the 1996 Guide for the Care and Use of Laboratory Animals (NIH) with approval from the Institutional Animal Care and Use Committee (IACUC) at Northeastern University. Pregnant female Sprague–Dawley rats were purchased from Charles River Laboratories (Wilmigton, MA) and arrived pregnant at embryonic day (E)15. All females were primiparous, and pregnant dams were housed singly. Parturition was checked daily, and the day of birth was denoted as P0. On P1, litters were culled to 10-12 pups, with 5-6 males and 5-6 females, when possible. Whole litters were randomly assigned to be reared under control (Con) or maternal separation (MS) rearing. Dams and pups were housed under standard laboratory conditions in polycarbonate wire-top caged with pine shave bedding and food and water available ad libitum. The facility was kept on a 12-h light/dark cycle (light period between 0700 and 1900) with regulated temperature (22–23°C) and humidity (37%–53%). All pups were toe clipped for identification on P5. Con pups were otherwise left undisturbed except for handling on P12 and P15 and testing days (P9, 10, 19, 20). All pups were weighed on P9 and P20.

### Maternal separation

MS was conducted daily from P2-20, during which pups were isolated from their dam and littermates with home cage bedding. From P2-P10, pups were placed in individual plastic cups kept to 37°C in a water bath for 3.5 h (0930–1300 h). From P11-P20 when pups were no longer poikilothermic, pups were placed in individual mouse cages for 4h. During the separation period, MS dams remained in their home cage in a separate room. After the separation period, pups were returned to the home cage with the dam and left undisturbed until the following day. Control litters remained with the dam and were handled by an experimenter during routine weighing (on P9 and P11) and cage-changes (2x/week).

### Maternal Behavior in Response to Pup USVs

Dam behavior in response to pup vocalizations impacted by adversity exposure were tested at two timepoints (Experimental Design illustrated in Figure 1). The dams were tested when pups were P10 and again at P20–ages at which pup vocalizations vary depending on MS or Con rearing. Different pups were used for the P10 and P20 tests.

**Figure 1.**
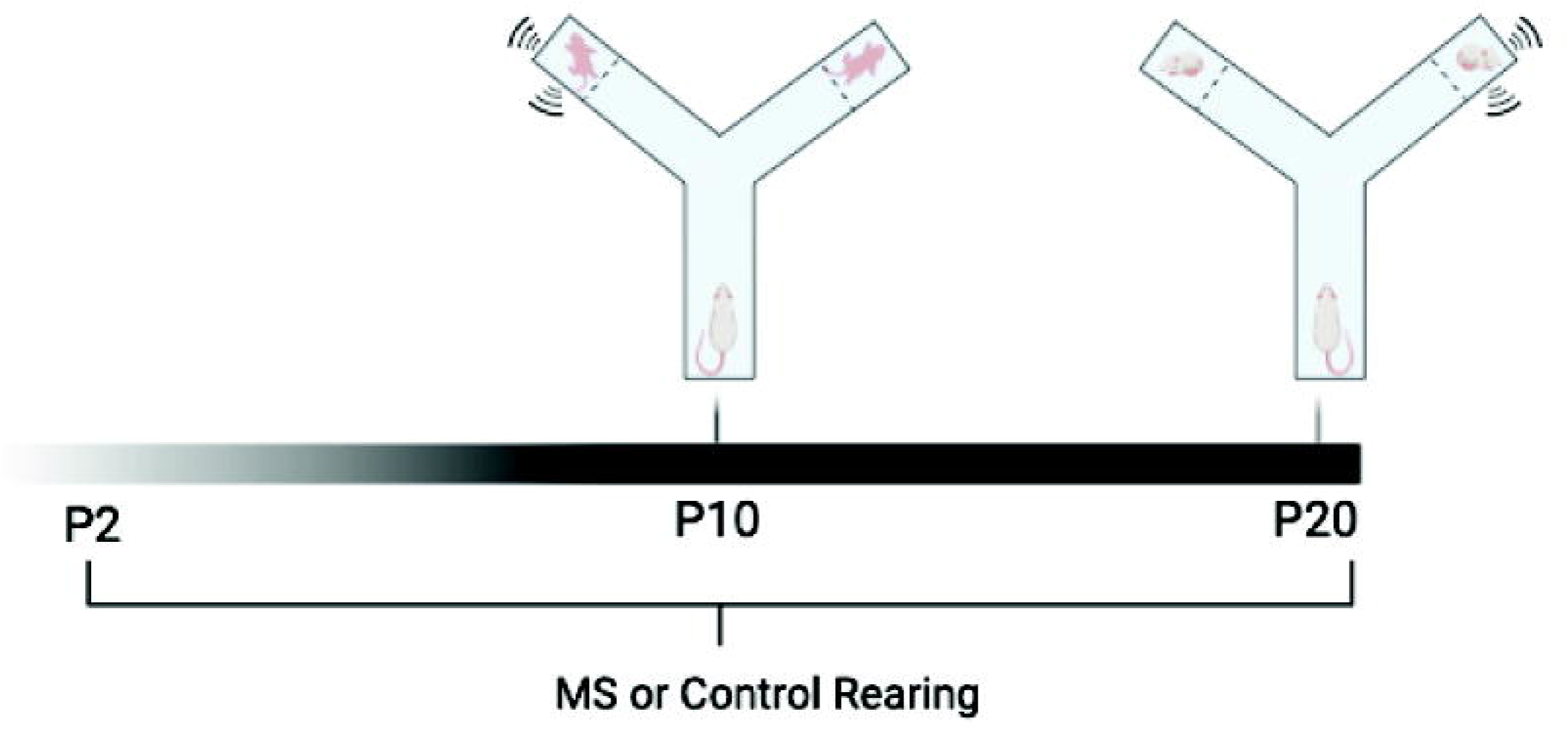
Experimental Design

A Y-maze apparatus was used to determine whether dams had preference for vocalizing pups over non-vocalizing pups and whether there was a sex bias. Dams were first allowed 5 minutes to habituate to the behavior room 24 h prior to testing. On the test day, one end of the Y-maze contained an anesthetized pup (to prevent vocalizations) on a weighboat. Pups were anesthetized with a ketamine/xylazine cocktail at a dose of 1mg/kg immediately prior to testing, and kept on a heating pad until testing to ensure that body temperature was not different between pups. A conscious, vocalizing pup of the same sex and age was placed on a weighboat in a second arm. A fine mesh barrier was placed on each of the two ends with pups to prevent retrieval and visual identification by the dam. The arms containing the vocalizing and non-vocalizing pups were counterbalanced to prevent arm bias. The dam was placed on the third arm of the Y-maze apparatus and behavior was recorded for a 5-minute period. The test was repeated with one male and one female pup pair for each dam, totaling two trials per dam. EthoVision video analysis software (Noldus) was used to analyze whether dams preferentially spend more time in the arm of a vocalizing pup over a non-vocalizing pup, and whether preference and latency to enter each arm was impacted by the sex of the pup or their adversity history. Outcome measures were the duration of time spent in the center of the Y-maze (i.e., no choice), frequency of entries into each arm, the latency to enter each arm, duration of time in each arm, and the percent time spent in each arm relative to time spent in all three arms.

### Ultrasonic Vocalization Recordings and Analysis

An ultrasonic microphone (Avisoft Bioacoustics, model CM16/CMPA) was used to validate the presence and absence of USV emissions from awake or anesthetized pups, respectively. The microphone was positioned 5 cm above the arm containing the vocalizing pup and recorded each pup for the 5-minute testing period. Audio files were uploaded and analyzed using DeepSqueak, a Matlab program for USV analysis. USVs were visualized by converting each file to spectrograms using Fast Fourier Transformation. Each call detected by Deepsqueak was manually confirmed or rejected by a trained experimenter (For more detailed methodologies, see Granata et al., 2021).

### Data Analysis

Group sizes were determined by power analyses with G*Power using effect sizes from previous maternal behavior studies, for a power of 0.8 for Y-maze analyses. Therefore, 8-10 dams/rearing conditions at each age were tested. Total time spent in each arm, as well as latency to enter and frequency of entries to each arm of the Y-maze were analyzed with Rearing x Pup Sex x Pup Condition (awake or anesthetized) mixed 3-way ANOVAs, with pup condition as a repeated measure. Time spent in the center of the Y-maze (i.e., time spent not making a choice) was analyzed with a Rearing x Sex 2-way ANOVA. Significant effects in 2- and 3-way ANOVAs were followed up with post-hoc multiple comparisons with Šidák’s correction.

## Results

USV analyses confirmed that awake pups emitted USV during the 5-min Y-maze task (523 calls ± 65 at P10; 59 calls ± 20 at P20), while anesthetized pups were silent. When litters were P10, a 3-way Sex x Rearing x Pup Condition ANOVA revealed that all dams spent a higher total time in the arm of the awake pup (main effect of Pup Condition *F*_1,28_ = 15.03; *p* = 0.0006; η^2^ = 0.219). A Rearing x Pup Condition interaction (*F*_1,27_ = 6.221; *p* = 0.019; η^2^ = 0.075) was found, and this effect was driven by a difference in which only MS dams presented with a choice between two male pups spent more time in the arm of the awake pup (*p* = 0.015) (Figure 2a). No significant differences (p > 0.05) were observed in the frequency of entries into each arm, the latency to enter each arm, or the time spent in the center of the maze (Figure 3a). General locomotion was also not different between groups, indicated by no differences in the total number of entries into all three arms (Figure 3b).

**Figure 2.**
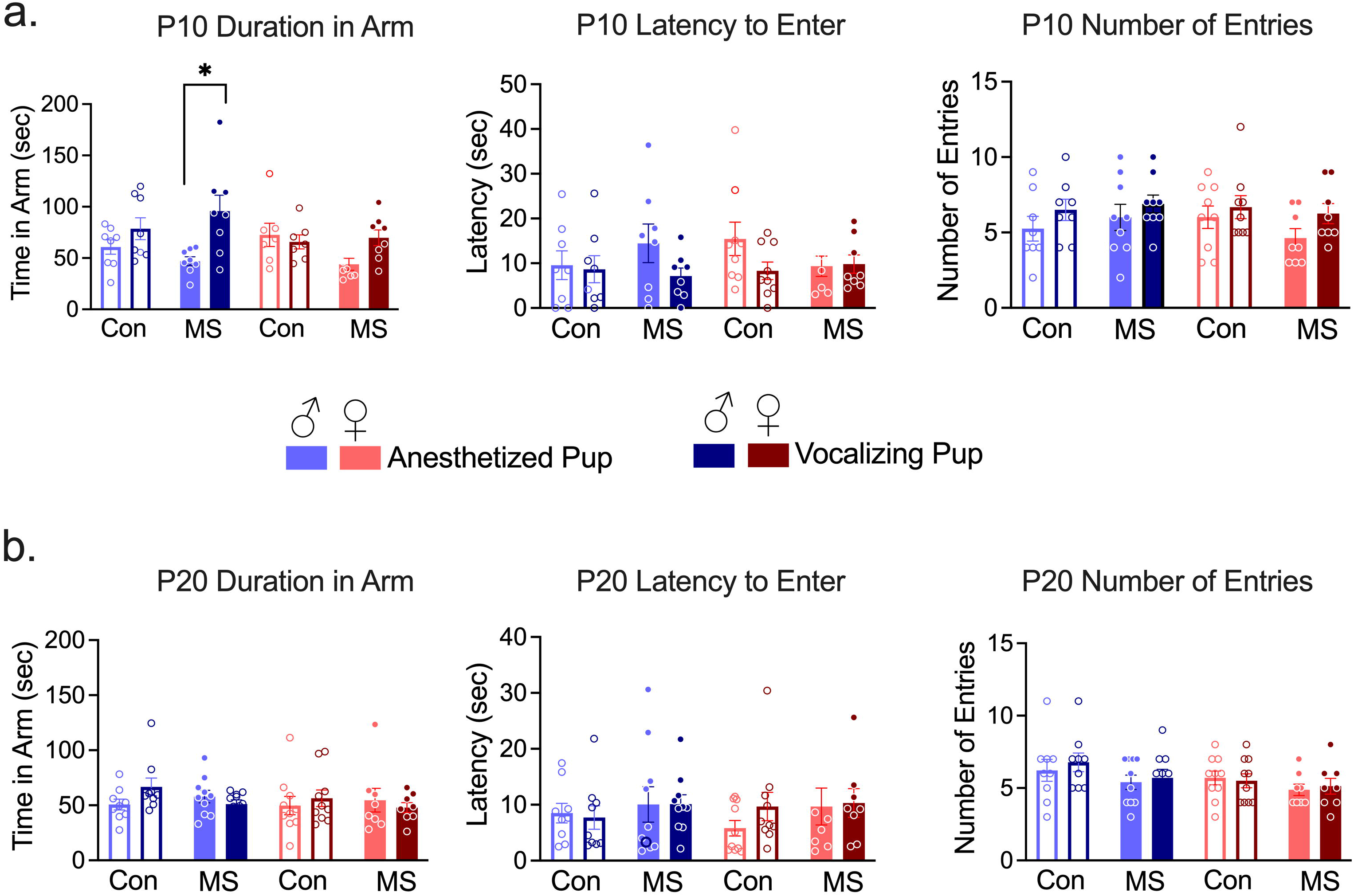
Maternal approach behaviors towards anesthetized or vocalizing pups within the Y-maze on (a) postnatal day (P)10, or (b) P20. Means ± SEM with individual data points are shown. *p < 0.05 difference between anesthetized and vocalizing pup, after correction for multiple comparisons. n = 8-10

**Figure 3.**
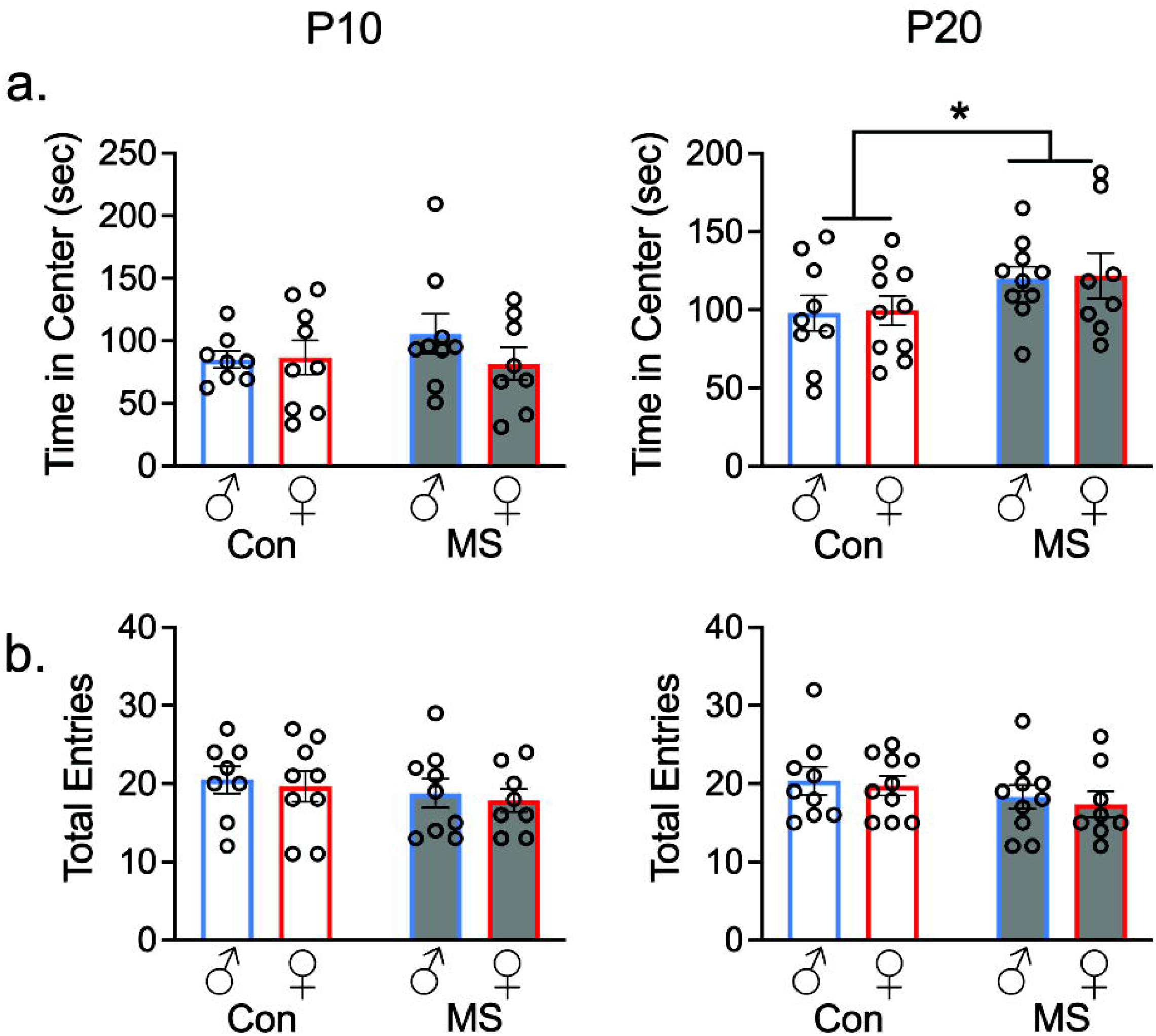
Time spent in the center of the Y-maze (a) and total locomotion (b) at P10 and P20. Means ± SEM with individual data points are shown. *p < 0.05 difference between MS and Con dams. n = 8-10

When litters were P20, no significant differences (p > 0.05) were found in a Sex x Rearing x Pup Condition 3-way ANOVAs on arm preference as assessed by duration spent in each arm, latency to enter each arm, or frequency of arm entries (Figure 2b). However, a 2-way Sex x Rearing ANOVA revealed that MS dams spent more time in the center of the Y-maze, compared to control dams (main effect of rearing *F*_1,33_ = 4.269; *p* = 0.047; η^2^ = 0.114) (Figure 3a). General locomotion was once again not different between groups, indicated by no differences in the total number of crosses between arms (Figure 3b).

## Discussion

The aim of this study was to determine how ELA affects pup-driven maternal behavior. We tested the influence of rearing environment on dam preference for vocalizing vs. non vocalizing pups by examining maternal interaction time, latency to approach, and arm entries. In order to reveal whether sex-specific consequences of ELA in offspring may be due to differences in maternal responsiveness to male versus female pups we compared responses to males and females at two pup developmental stages. We hypothesized that dams would have increased interaction times with vocalizing pups as compared to non-vocalizing pups, and performed an exploratory analysis on the impact of MS and pup sex on dam behavior. Here, we chose to investigate USV-driven maternal approach to expand upon prior findings from our lab and others showing that ELA alters predictability (Granata et al., 2022) and quality (Gallo et al., 2019) of maternal care, as well as USV communication (Granata et al, 2021).

We found that dams spent more time with vocalizing pups compared to non-vocalizing pups at the P10 time point, with a Rearing x Pup Condition interaction revealing that this difference was driven by MS. Notably, dams actively undergoing MS spent more time with male vocalizing pups as compared to male non-vocalizing pups, specifically. This is in line with prior work from our group showing that MS exposure exacerbated the age-related increase of total USV emissions by P10 in males only. Therefore, increased USV emissions may direct maternal care toward MS males. These effects are important to consider in light of myriad findings from our lab and others showing sex specific effects of MS exposure across all developmental stages (Granata et al., 2024; Granata et al., 2022b; Honeycutt et al., 2020). Our findings support the hypothesis that sex-specific pup-dam interactions may drive later life outcomes following adversity.

Importantly, we did not observe increased approach to vocalizing pups at P20, which may be associated with the relative lack of USV emissions by pups at this age. However, the finding that MS-exposed dams spent more time in the center of the Y-Maze may indicate that exposure to MS disrupts processes required for deciding between vocalizing and non-vocalizing pups.

## Conclusions

Here, we found that pup USVs drove dam-pup interaction time in a Y-Maze apparatus. This was affected by MS exposure in a sex specific manner, such that MS-exposed dams interacted with conscious male pups more than unconscious male pups. USV impacted dam approach on P10, but not P20, indicating that pup-driven maternal care is age specific. These findings illuminate the need to study sex dependent actions such as USVs and their interaction with maternal care within the home cage environment as predictors for sex specific outcomes following adverse rearing.

